# Neural Evidence That Gender-Inclusive Language Attenuates Automatic Gender Predictions

**DOI:** 10.64898/2026.07.16.738926

**Authors:** Luana Serafini, Martina Abbondanza, Francesca Pesciarelli

**Author notes:** These authors share first authorship (Luana Serafini, Martina Abbondanza).

## Abstract

Languages vary in whether and how they encode gender. As societies pursue greater inclusiveness, gender-inclusive language (GIL) has emerged as a debated innovation, introducing pronouns and morphological strategies to neutralize grammatical gender (e.g., *todxs*, for *todos/todas*, “all”). Yet whether GIL actually reduces gender expectations remains empirically underexplored. Evidence at the neural level is especially scarce. To address this gap, we asked Italian speakers to categorize masculine (*lui*, “he”) and feminine (*lei*, “she”) pronouns following nouns with canonical (masculine -*o*, feminine -*a*) or neutralized endings (-*\**, -*ə*). Noun gender-stereotypical association was independently manipulated (male-oriented, e.g., *chirurgo/a/ǝ/\**, “surgeon”; female-oriented, e.g., *maestra/o/ǝ/\**, “teacher”; neutral, e.g., *vicina/o/ǝ/\**, “neighbor”). Recording response times and event-related potentials (ERPs) to target pronouns allowed us to assess their automatic integration and processing costs. Gender-incongruent pronouns after canonical forms elicited longer response times and larger N400 and P300 amplitudes, reflecting increased processing difficulty and reanalysis. Pronouns following GIL forms showed an intermediate processing profile. Response times fell between consistent and inconsistent conditions, N400 amplitude was larger than in the consistent condition, and P300 amplitude was reduced compared to the inconsistent condition. Together these findings indicate that both pronouns remained viable interpretations and could ultimately be integrated. Stereotype-based knowledge shaped early processing without preventing integration, predominantly for masculine pronouns. Overall, our findings provide the first neural evidence that, despite processing costs, GIL forms *can* foster gender-neutral representations, with relevance to ongoing societal debates and broader cross-linguistic implications.

## Main Text

Inclusiveness is central to achieving equality and accounting for diversity in our society. Language, as a dynamic social tool, adapts to societal demands. The emergence of gender-inclusive language (GIL) exemplifies this process, providing novel means to refer to individuals whose identities—including non-binary and gender-diverse categories—are not captured by traditional male–female dichotomies (Sczesny et al., 2016). Existing language systems can be broadly divided into three main categories based on how gender is expressed: genderless languages, natural gender languages, and grammatical gender languages (Corbett, 1991). Genderless languages, e.g., Turkish and Finnish, do not grammatically convey gender; Natural gender languages, e.g., English, convey gender mainly through pronouns (e.g., masculine “he”, feminine “she”); Grammatical gender languages, e.g., French, Spanish, Italian, and German, pervasively encode gender via morphological marking (e.g., *maestr-**a**/maestr-**o***, “female/male teacher”) and through agreement processes affecting nouns, determiners, adjectives, and verbs (e.g., *l**a** maestr**a** brav**a** è tornat**a*** “the good (female) teacher is back”).

To generically refer to an individual whose gender is unknown or to a mixed-gender group, the generic masculine is an option—that is, masculine forms used to refer to individuals without highlighting their gender. Such forms appear to elicit male-biased mental representations in both natural gender languages (e.g., *mankind* used to denote humankind) and grammatical gender languages (e.g., the Italian masculine ending *-o* in *medico* “doctor,” used generically) (Glim et al., 2024; Gygax et al., 2008; Misersky et al., 2019;), thereby failing to represent non-male individuals. Besides grammar, gender is also conveyed through stereotypes—shared beliefs about how women and men are or should be (Ellemers, 2018). Thus, even grammatically neutral role nouns when stereotypically associated with men or women (e.g., *doctor, nurse*), can elicit gendered expectations, implicitly and automatically (Pesciarelli et al., 2019; Proverbio et al., 2018; Siyanova-Chanturia et al., 2012), thus limiting inclusiveness.

In recent years, the need for gender inclusiveness has prompted bottom-up linguistic innovation. In natural gender languages, emphasis falls on neutral pronouns like singular *they* in English or neologisms such as *hen* in Swedish (Arnold et al., 2021; Morris et al., 2026; Vergoossen et al., 2020). Grammatical gender languages innovate morphologically to neutralize endings: French employs the mid-dot (*citoyen·ne·s* “male and female citizens”; Pozniak et al., 2024; Tibblin et al., 2023), German the gender star (*Lehrer*innen* “male and female teachers”; Glim et al., 2025; Körner et al., 2022), while Spanish and Italian adopt innovative non-binary morphemes (e.g., Spanish: *-e/-x*: *todes/todxs* “all”; Román Irizarry et al., 2025; Italian: *-*/-ǝ*: *tutt*/tuttǝ* “all”; Abbondanza et al., 2025; Comandini, 2021; Mazzucca et al., 2026). In light of the societal urgency for GIL and the debates it generates (Sulis & Gheno, 2022; Coady, 2024), it is essential to establish empirical evidence regarding its capacity to foster gender-neutral representations, as well as the processing costs it may impose. Although evidence is accumulating rapidly, it remains relatively scarce. Focusing on innovative morphology, German gender star and French midpoint showed evidence for a female bias (Glim et al., 2025; Körner et al., 2022; Pozniak et al., 2024; Tibblin et al., 2023). Spanish and Italian non-binary endings showed elevated processing costs as compared to binary norms —shaped by lexical frequency, stereotypes, and diversity attitudes for Spanish non-binary endings— without compromising overall understanding (Marenghi et al., 2026; Román Irizarry et al., 2025; Stetie & Zunino, 2022; Zarwanitzer & Gelormini-Lezama, 2023). Abbondanza et al. (2025) showed that the Italian schwa attenuates, but does not eliminate, gendered expectations, with effects depending on lexical properties and presentation modality.

Neural measures can provide decisive evidence on both effectiveness and processing costs. Electroencephalography (EEG), particularly event-related potentials (ERPs), captures language processing at millisecond resolution, offering a fine-grained view of how novel morphological forms are integrated. To date, only one study has examined the neural representation of such forms, focusing on the German gender star. This study reported a P600 effect for masculine continuations (Glim et al., 2025)—an ERP component typically associated with syntactic reanalysis or morphosyntactic repair (Friederici, 2011)—suggesting that masculine continuations are processed as violations.

Neural evidence is still needed to determine whether gender-inclusive forms effectively neutralize grammatical gender cues and also attenuate gender stereotypes. We investigate these questions in Italian using a priming paradigm. Because this paradigm has been extensively validated in the assessment of the automatic and implicit activation of grammatical and stereotypical gender expectations (Banaji & Hardin, 1996; Casado et al., 2023, 2026; Chalyvidou & Weber, 2025; Pesciarelli et al., 2019; Siyanova-Chanturia et al., 2012), it provides a solid foundation for directly comparing automatic gender expectations elicited by canonical forms versus forms manipulated with inclusive suffixes. This approach typically presents a prime (e.g., a grammatically gender-marked or stereotypically biased word) followed by a target (e.g., third-person pronouns such as *he* or *she*). Grammatical prime–target mismatches reliably elicit N400 effects (200–400 ms; indexing semantic integration difficulty; Kutas & Federmeier, 2000) as well as later P300 effects (400–550 ms; associated with syntactic repair similarly to the P600, or context updating; Donchin & Coles, 1988), whereas stereotypical mismatches selectively elicit N400 effects. We adapted the paradigm by pairing Italian pronoun targets (*lui* “he”, *lei* “she”) with primes in canonical (masculine *-o*, feminine*-a*) or gender-inclusive (*-\**, *-ǝ*) forms, varying in stereotype associations (neutral, male-oriented, female-oriented). Asterisk and schwa forms (e.g., male-associated: *avvocat*/avvocatǝ* “lawyer”) were chosen as they dominate as primary gender-inclusive strategies in Italian (Comandini, 2021). This allowed us to test whether inclusive forms support gender-neutral representations, indexed by equal integration of masculine and feminine targets. We first test whether neutralization eliminates morphology-driven gender expectations, predicting mismatch effects (N400, P300) for *-o/lei* and*-a/lui*, but no such mismatch effects for *-*/lui, -*/lei or -ǝ/lui, -ǝ/lei* if these effectively neutralize gender predictions. We then assess stereotype-driven influences, expecting stronger effects when morphology and stereotypes both mismatch the pronoun (e.g., *avvocat**o*** “male lawyer”- *lei* “she”), and potential residual stereotype effects even for neutral forms (e.g., *avvocat\** “lawyer”- *lei* “she”).

Gender-inclusive language has become the focus of intense public, political, and academic debate, yet little is known about how these novel linguistic forms are processed in real time. This work fills a critical gap in research on gender-inclusive forms, offering neural evidence that neutralized word endings can indeed foster gender-neutral representations, wherein both masculine and feminine genders can be successfully integrated, largely irrespective of gender stereotypes. The paradigm is generalizable across languages and provides a framework to study how the mind adapts to linguistic change.

## Methods

### Participants

Thirty-nine monolingual native Italian speakers participated in the experiment (21 women, age range = 18 - 31 yrs, *M* = 22.54 yrs, *SD* = 2.63 yrs). All participants had normal or corrected-to-normal vision and were right-handed, with the exception of one left-handed, as assessed by an Italian version of the Edinburgh Handedness Inventory (Oldfield, 1971). None of the participants reported a history of neurological or language-related disorders. Three participants were excluded from the analyses: one who reported having dyslexia, one due to illness during the session and one due to excessive EEG artifacts. The final sample consisted of 36 participants (20 women, age range = 18-31 yrs, *M* = 22.47 yrs, *SD* = 2.71 yrs), in line with previous literature (Pesciarelli et al., 2019; Siyanova-Chanturia et al., 2012). The experiment was approved by the local Ethical Committee CEAR (Comitato Etico per la Ricerca UniMoRe) and it was run in accordance with the “Italian Association of Psychology” (AIP) Ethical Guidelines (Codice Etico: https://aipass.org/chi-siamo/#ethical-code) and the Declaration of Helsinki. All participants gave written informed consent, and university students received course credits.

### Stimuli

A set of 110 Italian words referring to professions, social roles, or personal characteristics served as prime stimuli. The items consisted of nouns and adjectives and were presented in four morphologically inflected forms: the canonical feminine form *-a* (e.g., *vicin-a* [G,F], “female neighbor”), the canonical masculine form *-o* (e.g., *vicin-o* [G,M], “male neighbor”), and two gender-neutralized forms: *-ə* (e.g., *vicin-ə* [G,N], “neighbor”) and *-\** (e.g., *vicin-\** [G,N], “neighbor”).

The stimuli were selected to represent three stereotypical gender categories. Thirty items had no stereotypical gender association (S,N; e.g., *vicina/o/ə/\**, “neighbor”); 30 were stereotypically associated with men (S,M; e.g., *chirurgo/a/ə/\**, “surgeon”); and 30 were stereotypically associated with women (S,F; e.g., *maestra/o/ə/\**, “teacher”). This design allowed morphological endings to be manipulated independently of stereotype content and stereotype content to be examined across morphological forms, yielding 12 combinations of grammatical gender marking and stereotypical association (see Figure 1b). An additional set of 20 common-gender nouns with no stereotypical gender association (e.g., *ospite* [G,M/F], “guest”) served as fillers. The 90 experimental items and 20 fillers were selected based on questionnaire data (see Supplementary Material).

**Figure 1.**
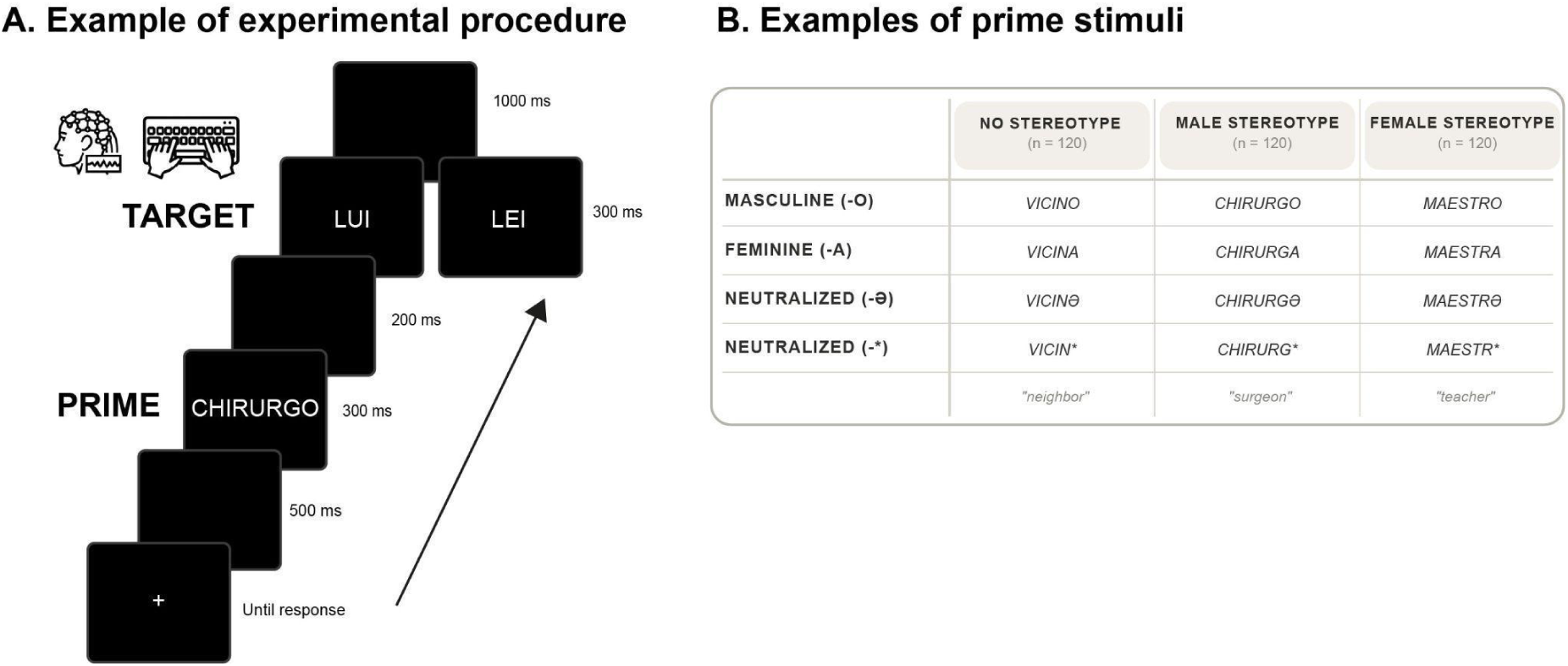
Task design and stimulus examples. (A) Schematic trial structure. Each trial begins with the presentation of a prime stimulus, followed by a target stimulus. Participants indicated whether the target was masculine or feminine. EEG was continuously recorded throughout the task. (B) Examples of prime stimuli. Prime stimuli consisted of Italian gender-marked words ending in canonical masculine or feminine suffixes or in neutralized forms. Words were classified into three categories based on stereotypical association: neutral, male-stereotypical, and female-stereotypical.

Experimental stimuli did not differ in lexical frequency (itTenTen20 corpus; Jakubíček et al., 2013), length, or valence, and the two stereotypical conditions did not differ in stereotype strength (Table S1 in Supplementary Material).

The selected words served as primes for two target stimuli: the Italian third-person singular pronouns *lui* [G,M] (“he”) and *lei* [G,F] (“she”). Each prime was paired with both pronouns, generating grammatically congruent and incongruent conditions, and gender-inclusive conditions, within each stereotype category (Table 1), as well as stereotypically congruent and incongruent conditions within each grammatical category (Table 2).

**Table 1.**
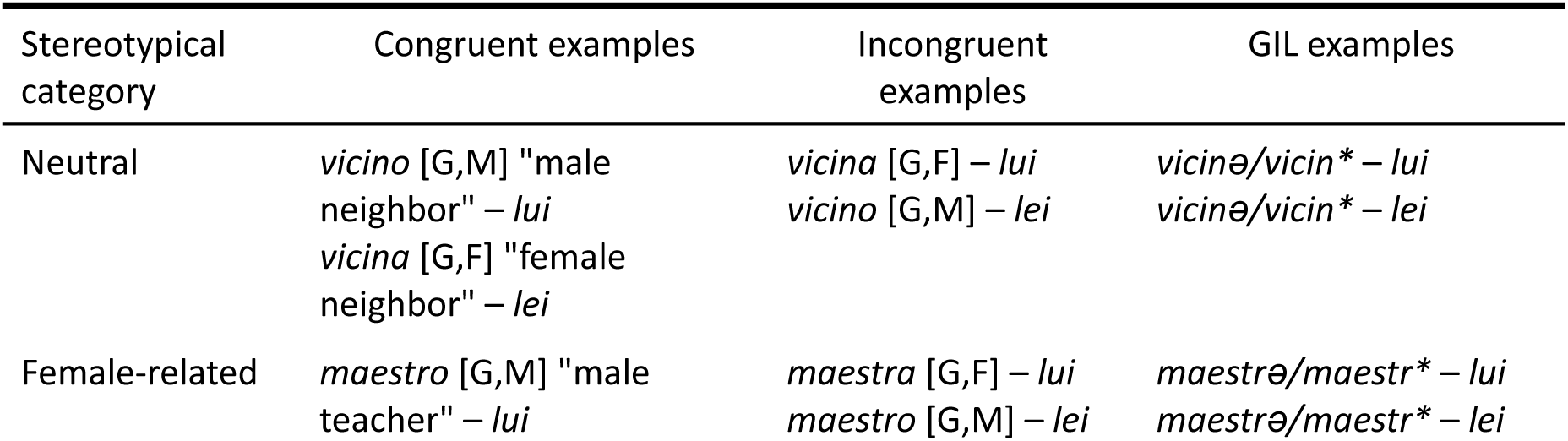
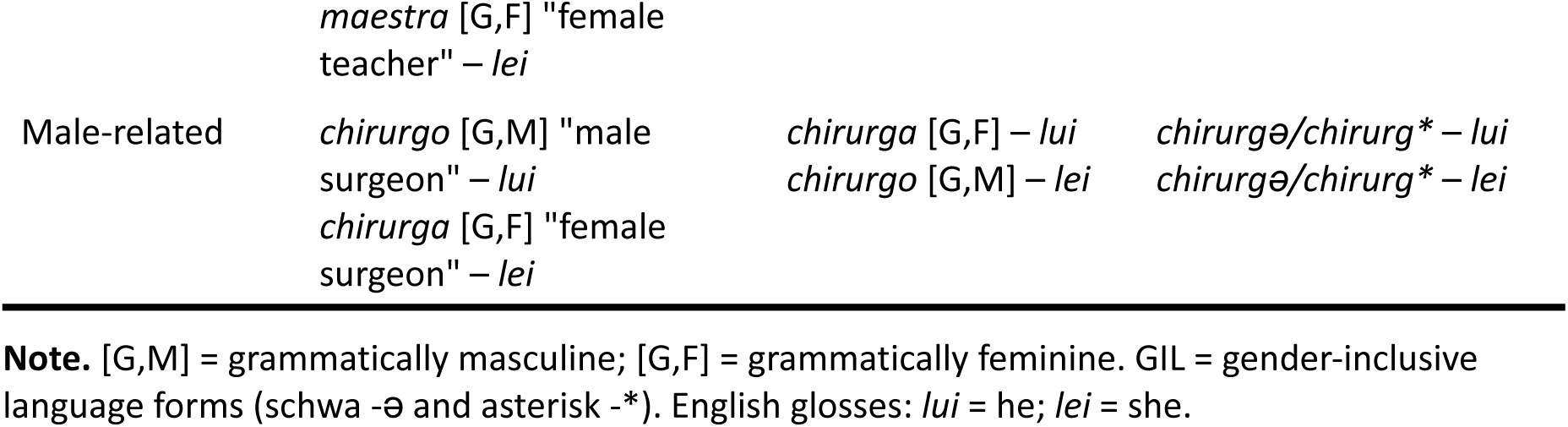
Examples of grammatically gender congruent, incongruent and gender-inclusive prime–pronoun pairs by stereotypical category.

**Table 2.**
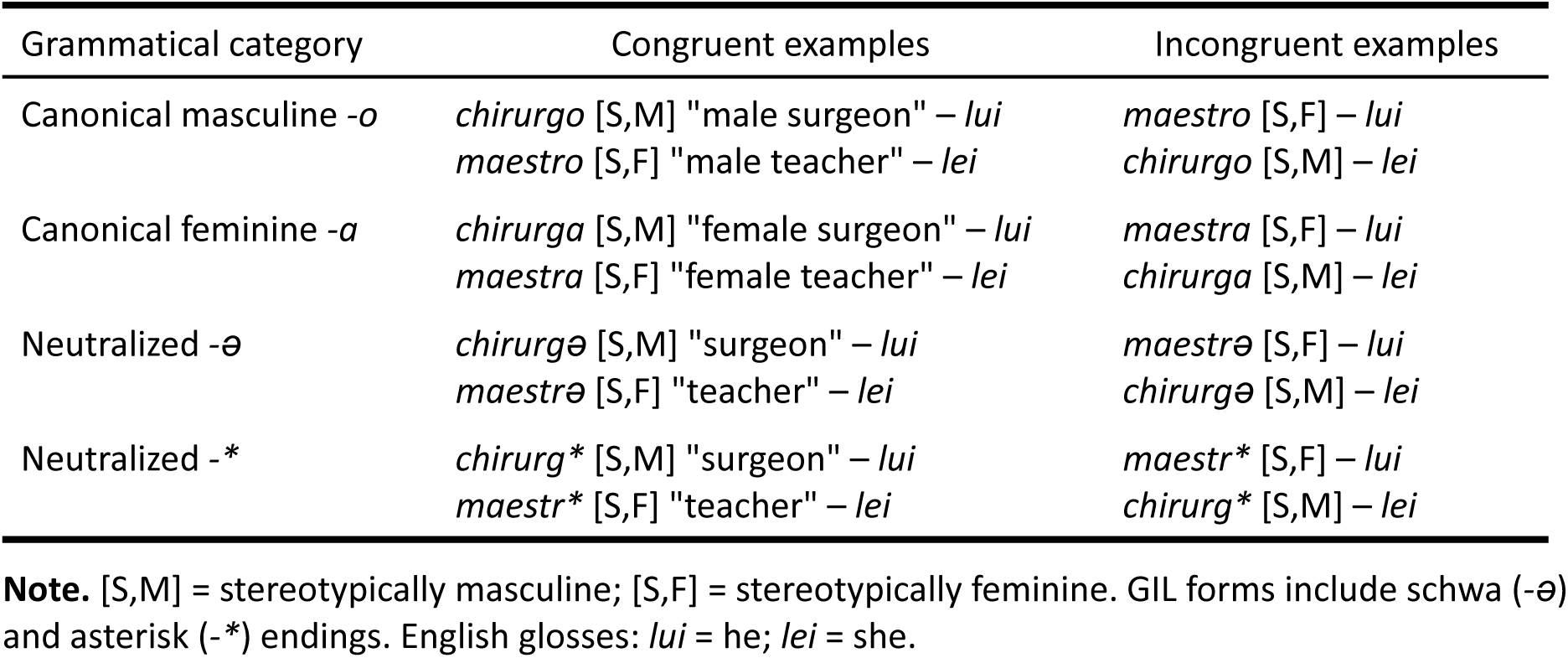
Examples of stereotypically gender congruent and incongruent prime–pronoun pairs by grammatical category.

### Design and Procedure

Participants were seated in a darkened, electrically shielded and sound-attenuated room. The experiment was implemented with the E-Prime software (Psychology Software Tools, 2016). Stimuli were presented at the center of a 24-inch LCD monitor (1920 × 1080 resolution, 59.94 Hz refresh rate), positioned approximately 70 cm from the participant at eye level. Prime words subtended 1.2–3.5° × 0.5° visual angle, whereas target words subtended 1.13° × 0.63° visual angle. Stimulus presentation was synchronized with the monitor refresh rate. Stimuli were displayed in white letters (Arial font, primes: 17 pt, targets: 22 pt) against a black background. Priority settings were optimized to ensure accurate display durations.

The procedure was adapted from Siyanova-Chanturia et al. (2012). Each trial, presented in randomized order, consisted of the following events: in the center of the monitor, a fixation cross (+) appeared until the participant’s response. Next, a blank screen appeared for 500 ms, then the prime word was displayed on the screen for 300 ms, followed by a blank screen for 200 ms. Subsequently, the target pronoun (*lui* or *lei*) was presented for 300 ms followed by a blank screen of 1000 ms, during which the participant could respond (Figure 1a). Participants were instructed to attend to both primes and targets and to read the prime words silently. Their task was to categorize the target pronoun as masculine or feminine as quickly and accurately as possible. Responses were made with two response keys and key assignment was counterbalanced across participants.

Trials were organized into eight blocks, with the order of blocks randomized across participants. Each nominal root (e.g., *vicin-*) appeared eight times across the experiment, once with each combination of ending and pronoun. Repetitions of the same nominal root were presented in different blocks. Before the experiment, participants completed a 10-trial practice block including only stereotype-neutral stimuli that were not repeated in the main experiment.

After EEG recording, participants completed the Ambivalent Sexism Inventory (ASI; Glick & Fiske, 1996), the Bem Sex-Role Inventory (BSRI; Bem, 1974), the Feeling Thermometer (Zavala-Rojas, 2014) measuring attitudes toward the LGBTQIA+ community, and rated, on a 7-point Likert scale, their familiarity with GIL and the degree to which they perceived it as useful and applicable. Overall, participants reported a medium-to-high familiarity with GIL (*M* = 5.58, *SD* = 1.33), and an overall neutral attitude towards it (*M* = 4.72, *SD* = 1.41). These measures were included to examine the influence of participants’ personality traits and knowledge and attitudes towards GIL on implicit linguistic processing. Results for these measures are reported in the Supplementary material (Table S2). All data were collected using REDCap (Harris et al., 2009).

### EEG Recording and Analysis

EEG was recorded using an ActiCHamp Plus system (Brain Products) from 64 active electrodes mounted in an ActiCap Slim and positioned according to the international 10–10 system. Three additional electrodes recorded electrooculogram (EOG) activity (placed at the outer canthi of both eyes and below the left eye). Data was recorded at a sampling rate of 1000 Hz with FCz as the online reference and FPz as ground.

Offline preprocessing was conducted using EEGLAB (Delorme & Makeig, 2004) and ERPLAB (Lopez-Calderon & Luck, 2014) running in MATLAB (The MathWorks Inc, 2021). Data were downsampled to 500 Hz and re-referenced to the average of the mastoid electrodes. EEG was band-pass filtered from 0.1 to 30 Hz. Noisy electrodes were interpolated using spherical spline interpolation. No more than two per participant were interpolated (3.13% of electrodes). Ocular artifacts were corrected using the FastICA algorithm. Continuous data were segmented into epochs from −200 to 800 ms relative to target onset and baseline-corrected using the −200 to 0 ms pre-stimulus interval. Epochs were rejected if the signal exceeded ±100 µV or showed a voltage change greater than 200 µV within a 200-ms window. Artifact rejection resulted in an average data loss of 1.69% (*SD* = 2.49%) among the remaining participants. Epochs associated with correct responses were averaged separately for the 24 experimental conditions (mean trials per condition = 28.06, *SD* = 1.29, range = 23.83–29.83).

### Statistical Analysis

Accuracy in the gender decision task was at ceiling (95%), thus it was not analyzed. Reaction times (RTs) shorter than 200 ms or exceeding the response deadline of 1300 ms were removed, resulting in the exclusion of less than 1% of the data. RTs (log-transformed) of correct responses were analyzed using a linear mixed-effects model with Prime Ending (-o, -a, -schwa, -asterisk), Target Gender (masculine, feminine), and Prime Stereotype (male-oriented, female-oriented, none) as fixed effects, and random intercepts for participants and items. Results showed that the three-way interaction did not modulate RTs, but a two-way interaction between Prime Ending and Target Gender significantly predicted RTs. A model comparison between the full model including the three-way interaction Prime Ending × Target Gender × Prime Stereotype and a reduced model (Prime Ending × Target Gender + Prime Stereotype) showed that the more complex model did not improve fit (χ²(14) = 13.02, *p* = .53). We therefore report results from the more parsimonious model including Prime Ending × Target Gender + Prime Stereotype as fixed effects.

EEG data was analyzed using the Mass Univariate ERP Toolbox (Groppe et al., 2011) in MATLAB. Priming effects (i.e., differences between ERPs for incongruent and congruent prime-target pairs) and effects due to GIL (i.e., differences between ERPs for gender-inclusive and congruent/incongruent prime-target pairs), were tested for both feminine and masculine target pronouns, using repeated measures two-tailed cluster mass permutation tests (Bullmore et al., 1999) using a family-wise alpha level of 0.05. All time points between 100 and 800 ms at 40 electrodes of interest were included in the test (i.e., 14,000 total comparisons) and any electrode within approximately 4.32 cm of one another were considered spatial neighbors. Repeated measures one-sample t-tests were performed for each comparison using the original data and 2500 random within-participant permutations of the data. For each permutation, all *t*-scores corresponding to uncorrected p-values of 0.05 or less were formed into clusters. The “mass” of each cluster was the sum of the *t*-scores in the cluster, and the most extreme cluster mass in each of the 2501 sets of tests was used to estimate the distribution of the null hypothesis. This cluster-based approach offers superior spatial and temporal resolution compared to traditional mean amplitude ANOVAs while controlling for multiple comparisons.

## Results

### Longer Response Times in Grammatically Incongruent Conditions, While GIL Conditions Differ Significantly From Both Congruent and Incongruent Conditions

The linear mixed-effects model including Prime Ending × Target Gender + Prime Stereotype as fixed effects resulted in a significant interaction between Prime Ending and Target Gender (Figure 2). Specifically, being Prime Ending = asterisk and Target = *lui* (“he”) the reference levels, the output of the model was the following: Prime Ending -o:Target Gender -f: Est. = .109, *t* = 16.51, *p* <.001; Prime Ending -a:Target Gender -f: Est. = .059, *t* = 8.91, *p* < .001; Prime Ending -ǝ:Target Gender -f: Est. = .056, *t* = 8.45, *p* < .001).

**Figure 2.**
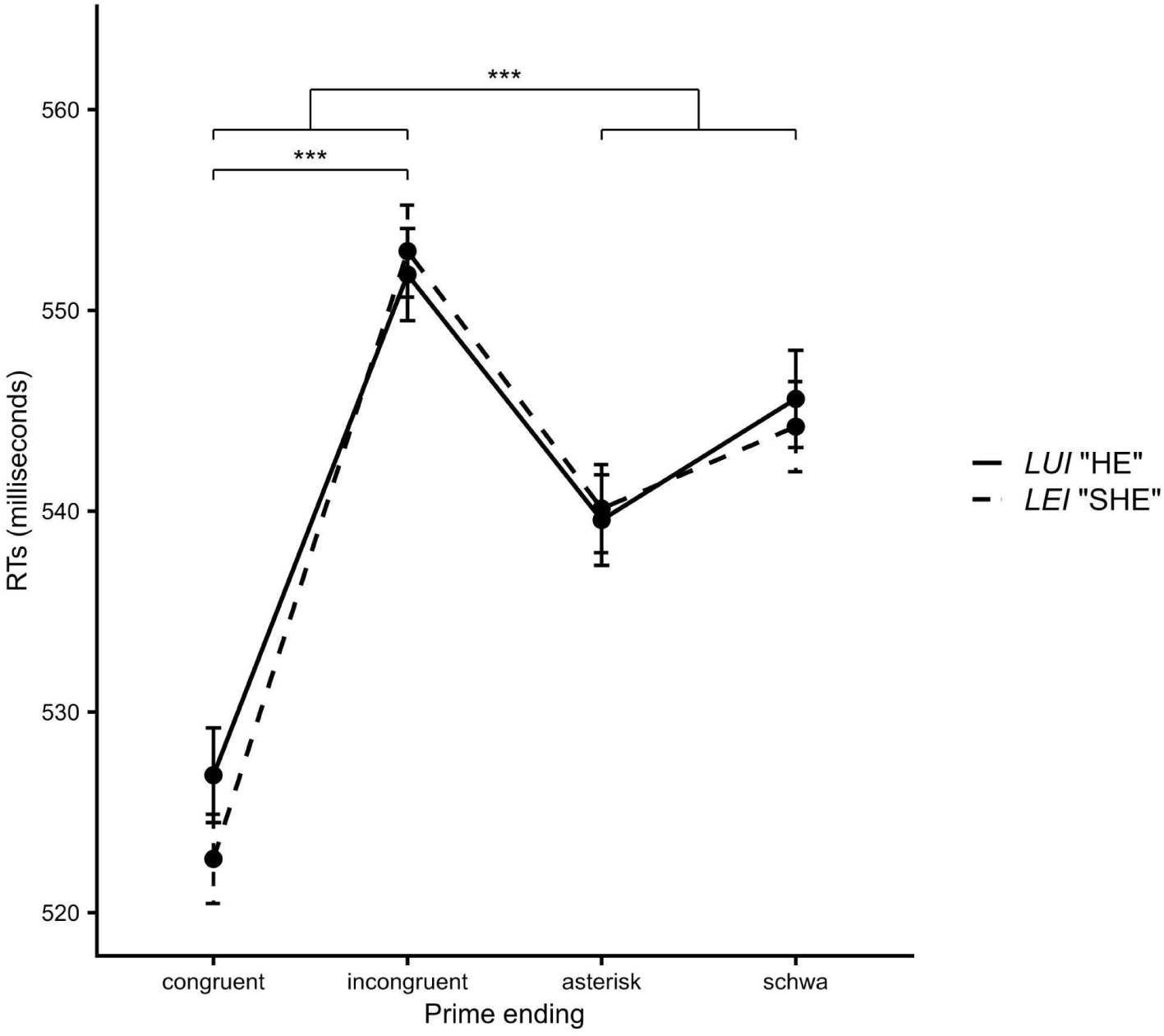
Longer response times in grammatically incongruent conditions, while GIL conditions differ significantly from both congruent and incongruent conditions. Average RTs when responding to the targets *lui* (“he”) and *lei* (“she”) based on the ending of the preceding noun (-a/-o/-*/-ǝ). Error bars represent standard errors of the means (SE).

Post hoc analyses (*emmeans*; Lenth & Piaskowski, 2026) showed that responses to the target *lui* were delayed when the preceding noun ended in *-a* compared with *-o* (*p* <.001), and responses to the target *lei* were slower when the preceding noun ended in *-o* compared with *-a* (*p* < .001). These results confirmed the gender-congruency effects that served as a baseline for investigating the GIL forms. Turning to the forms examined in the present study, RTs to the target *lui* following nouns neutralized with asterisk or schwa differed significantly from both the congruent and the incongruent conditions (*-a* vs. *-\**, *p* < .001; *-a* vs. *-ǝ*, *p* = .01; *-o* vs. *-\**, *p* < .001; *-o* vs. *-ǝ*, *p* < .001). The same pattern was observed for responses to the target *lei* (*-a* vs. *-\**, *p* < .001; *-a* vs. *-ǝ*, *p* < .001; *-o* vs. *-\**, *p* < .001; *-o* vs. *-ǝ*, *p* = .001), indicating that neither form influenced processing of the subsequent target by facilitating or inhibiting it. Interestingly, the two forms did not differ between each other across targets, showing that both asterisk and schwa symbols can be equally effective in neutralizing gender.

As a further step, we examined whether participants’ gender impacted on the reported effects, adding it in interaction with Prime Ending and Target Gender, obtaining a three-way interaction model. The random effect structure was identical to the main model. Results showed that participants’ gender did not influence the results (*p* > .05 for all the levels of comparison).

### Typical Grammatical Gender ERP Priming Effects, While GIL Conditions Differ Initially From the Congruent Condition and Later From the Incongruent Condition for Both Pronouns

First we examined the EEG response to pronouns primed by nouns in masculine (*-o*), feminine (*-a*), and gender-neutral forms (*-ə*, *-\**), irrespective of their stereotypical association, aiming to assess whether neutralization strategies effectively reduce morphologically triggered gender expectations more generally (see Figure 3).

**Figure 3.**
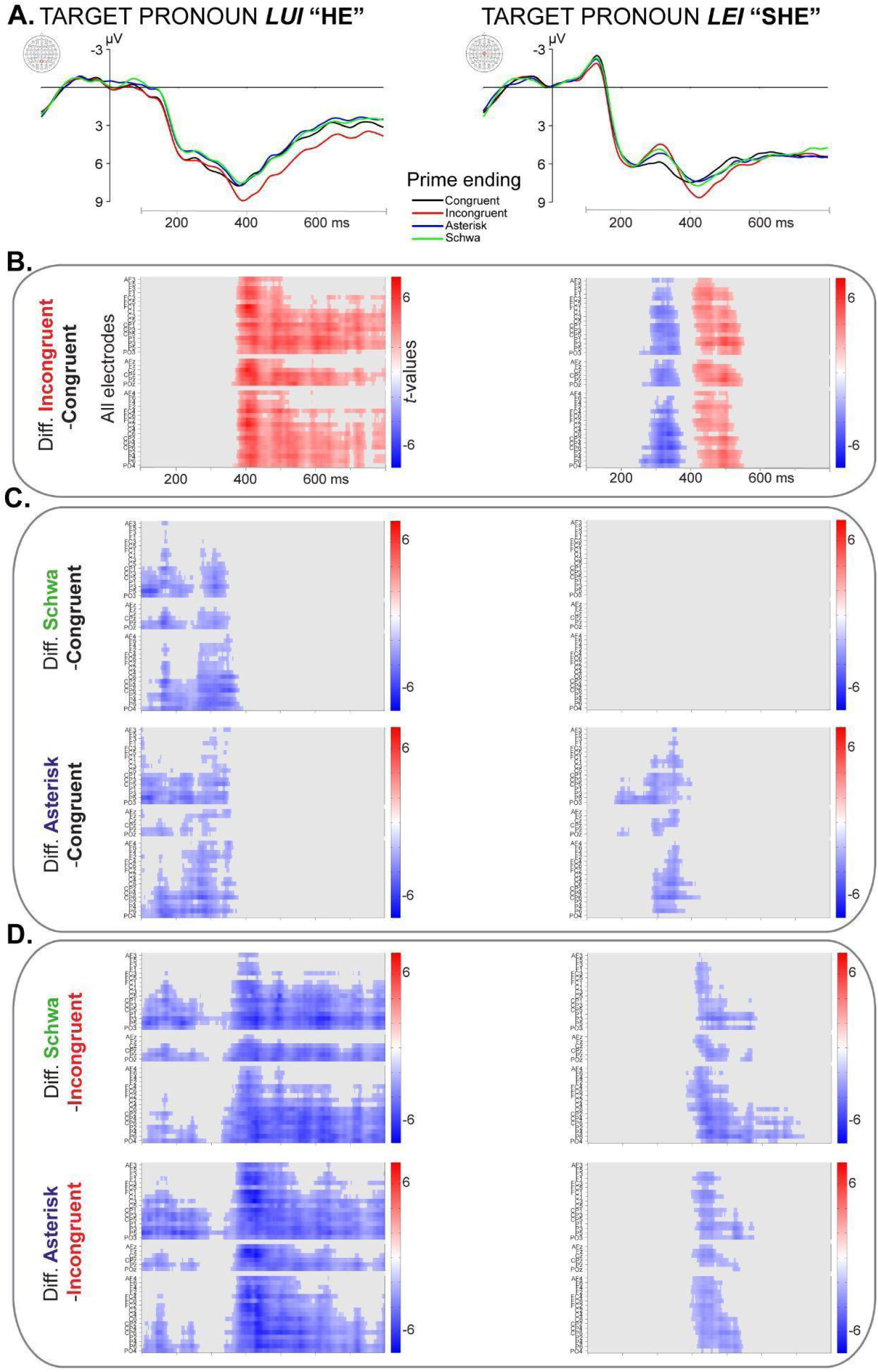
Typical grammatical gender ERP priming effects, while GIL conditions differ initially from the congruent condition and later from the incongruent condition for both pronouns. (A) EEG time-locked to target pronoun onset as a function of prime ending (color coded; data for EEG waveforms were low-pass filtered to 20Hz for visualization only). (B) Raster plots representing the results of the cluster mass permutation tests on EEG amplitude for target pronouns primed by incongruent as compared to congruent endings. (C) Raster plots representing the results of the cluster mass permutation tests on EEG amplitude for target pronouns primed by gender-inclusive as compared to congruent endings. (D) Raster plots representing the results of the cluster mass permutation tests on EEG amplitude for target pronouns primed by gender-inclusive as compared to incongruent endings. (B-D) Results are represented as a function of space (40 electrodes of interest) and time (100-800 milliseconds after target onset). Red indicates positive clusters, blue negative clusters.

In response to the masculine pronoun *lui* “he”, the grammatically gender-incongruent condition *-a*, compared with the congruent condition *-o*, yielded a statistically significant positive cluster (mass = 21207.545; *p* = 0). This effect extended from approximately 362 to 800 ms after pronoun onset, initially spanning all electrode sites and becoming more pronounced over centro-parietal sites from around 530 ms onward. Its latency and scalp distribution are consistent with a P300 priming effect extending into a later Late Positive Potential (LPP) effect. In response to the feminine pronoun *lei* “she”, the incongruent condition *-o*, compared with the congruent condition *-a*, yielded two statistically significant clusters. The first was a negative cluster (mass = −4847.005, *p* = .025) spanning all electrode sites from approximately 254 to 386 ms after pronoun onset. The second was a positive cluster (mass = 6634.567, *p* = .012), extending across all sites from approximately 406 to 554 ms. These effects are consistent, in both latency and scalp distribution, with N400-P300 priming effects (Figures 3a and 3b). For both pronouns, results show the expected grammatical gender priming effects, serving as a baseline for the investigation on inclusive forms.

We then tested how pronouns are processed following primes in gender-inclusive forms, compared with both grammatically gender-congruent and -incongruent primes (Figures 3c and 3d).

In response to the pronoun *lui*, we found that the GIL condition with primes ending in *-ə* relative to the congruent condition *-o*, yielded a statistically significant negative cluster (mass = −7438.183, *p* = .015), primarily over centro-parietal sites, extending from approximately 100 to 390 ms after pronoun onset. This pattern may reflect early unpredicted N100–lack-of-P200 effects, followed by an N400 effect. Compared with the incongruent condition *-a*, the same condition yielded a significant negative cluster (mass = −27392.729, *p* = 0), again maximal at centro-parietal sites and spanning the full 100–800 ms window, possibly reflecting an early N100-lack-of-P200 effect and a later sustained *absence* of P300–LPP effects. Similarly, the gender-inclusive condition with primes ending in *-\** differed significantly from both the congruent condition (negative cluster: mass = −7779.806, *p* = .013) and the incongruent condition (negative cluster: mass = −30661.509, *p* = 0), with comparable spatiotemporal characteristics to the schwa condition. Responses to *lui* did not differ between schwa and asterisk conditions (*p*s > .10). This pattern suggests that the masculine pronoun under gender-inclusive priming initially elicits integration difficulty (as reflected in early negativities) as compared to canonical grammatical forms, continuing as an N400 effect when compared to the congruent form, but is ultimately successfully integrated, as indicated by the absence of the P300–LPP effect observed in the incongruent condition.

In response to the pronoun *lei*, we found that the gender-inclusive condition with primes ending in *-ə* did not differ significantly from the congruent condition *-a*, but did differ from the incongruent condition *-o*, yielding a significant negative cluster (mass = −6285.800, *p* = .020). This effect extended from approximately 390 to 720 ms after pronoun onset, initially across all sites and later concentrating over right centro-parietal sites, consistent with the *absence* of a P300 effect. In contrast to the schwa condition, primes ending in *-\** also produced a significant difference from the congruent condition (negative cluster: mass = −3750.166, *p* = .045), spanning approximately 190 to 420 ms, with a distribution shifting from left parietal sites to all sites, consistent with an N400 effect. As with schwa, the asterisk condition also differed from the incongruent condition (negative cluster: mass = −4604.171, *p* = .032), with a shorter temporal extent (up to ∼590 ms), again consistent with the absence of a P300 effect. Responses to *lei* did not differ between schwa and asterisk conditions (*p*s > .10). Overall, this pattern suggests that the feminine pronoun under gender-inclusive priming shows a more limited early integration difficulty (no initial negativity, N400 effect restricted to the asterisk condition). As for the masculine pronoun, *lei* is also ultimately successfully integrated, as indicated by the absence of the P300 effect observed in the incongruent condition.

### Typical ERP Priming Effects Are Restricted to Stereotypical Primes; GIL Conditions Show Early Differences From Congruent for *Lui* and Later Differences From Incongruent Conditions, With Modulation by Stereotype for *Lei*

We then compared EEG responses to pronouns primed by words in the different endings, within each stereotype category (neutral, female-related, male-related) (see Table 1), to assess whether the priming and gender-inclusive effects depend on gender stereotype (Figure 4).

**Figure 4.**
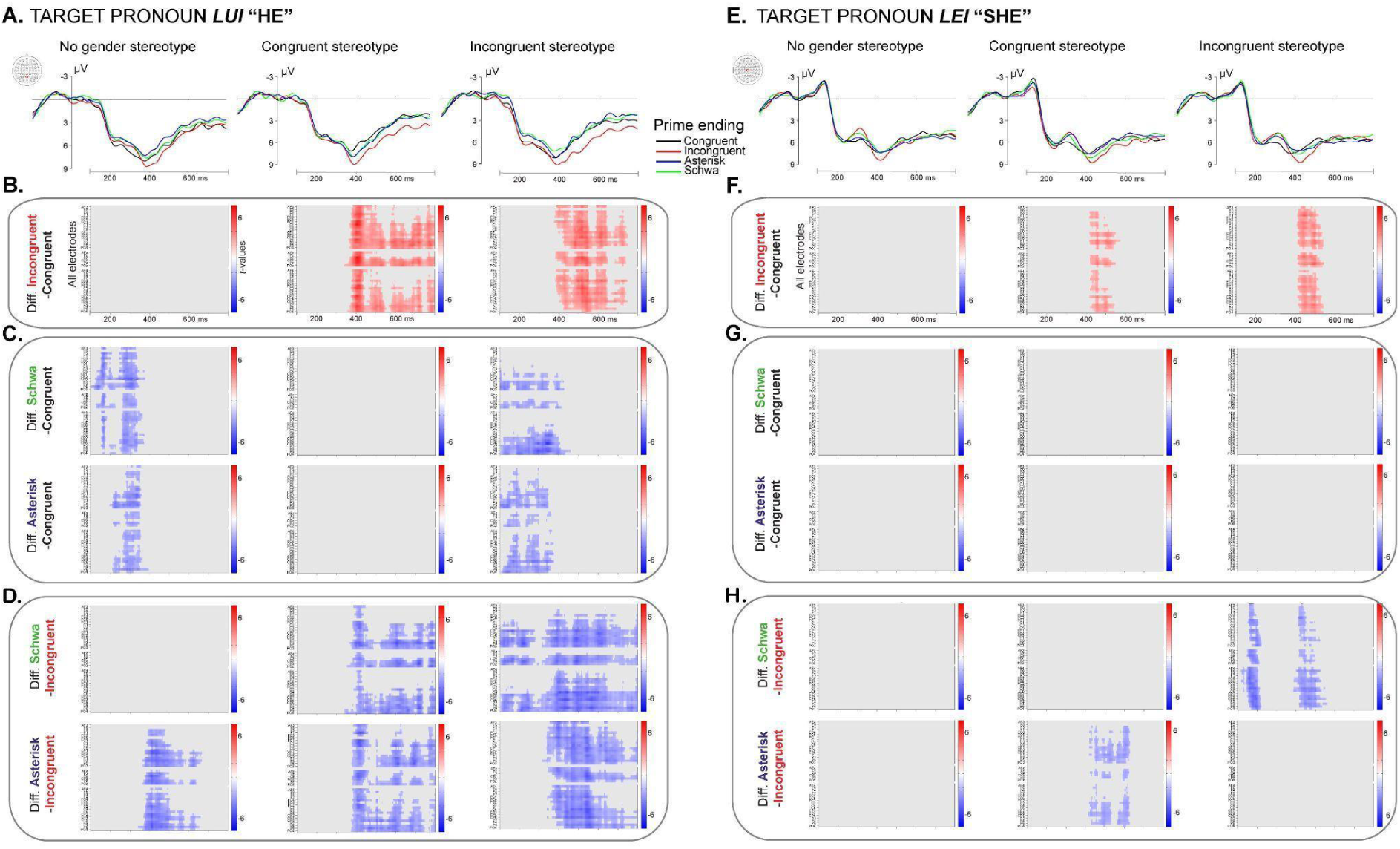
Typical ERP priming effects are restricted to stereotypical primes; GIL conditions show early differences from congruent for lui and later differences from incongruent conditions, with modulation by stereotype for lei. (A,E) EEG time-locked to target pronoun onset as a function of prime ending (color coded; data for EEG waveforms were low-pass filtered to 20Hz for visualization only) divided by stereotypical category. (B,F) Raster plots representing the results of the cluster mass permutation tests on EEG amplitude for target pronouns primed by incongruent as compared to congruent endings across stereotype categories. (C,G) Raster plots representing the results of the cluster mass permutation tests on EEG amplitude for target pronouns primed by gender-inclusive as compared to congruent endings across stereotype categories. (D,H) Raster plots representing the results of the cluster mass permutation tests on EEG amplitude for target pronouns primed by gender-inclusive as compared to incongruent endings across stereotype categories. (B-D) Results are represented as a function of space (40 electrodes of interest) and time (100-800 milliseconds after target onset). Red indicates positive clusters, blue negative clusters.

First we found that grammatical gender priming effects emerged only for stereotypically gender associated words. In response to *lui* “he”, the grammatically gender-incongruent condition *-a*, compared with the congruent condition *-o*, yielded a significant effects for stereotypically gender-congruent words (e.g., *chirurga*/*o*, “female/male surgeon”), expressed as a positive cluster (mass = 17152.864, *p* = 0) extending from approximately 342 to 800 ms after pronoun onset, initially across all sites and later concentrating over centro-parietal sites (from ∼450 ms); and for stereotypically gender-incongruent words (e.g., *maestra/o*, “female/male teacher”), with a positive cluster (mass = 14781.559, *p* = .0008) spanning approximately 376 to 748 ms across all sites. In response to *lei “she”*, the grammatically gender-incongruent condition *-o*, compared with the congruent condition *-a*, yielded significant effects for stereotypically gender-congruent words, expressed as a positive cluster (mass = 3338.619, *p* = .039) extending from approximately 416 to 572 ms mostly at centro-parietal sites; and for stereotypically gender-incongruent words, with a positive cluster (mass = 5719.235, *p* = .013) spanning approximately 410 to 548 ms across all sites. No grammatical gender priming effect emerged for stereotype-neutral words (e.g., *vicina/o*, “female/male neighbor”) for either pronouns (*p*s > .06). These results suggest that gender stereotypes contribute additively to grammatical gender priming effects for both masculine and feminine pronouns.

We then examined how pronouns are processed following gender-inclusive primes, compared with both congruent and incongruent grammatical gender primes, within each stereotype category.

In response to the pronoun *lui*, (i) for stereotype-neutral words (e.g., *vicina/o*, “female/male neighbor”), both *-ə* and *-\** conditions differed significantly from the congruent condition *-o*, yielding negative clusters (schwa: mass = −6976.400, *p* = .013; asterisk: mass = −4573.749, *p* = .034) extending from approximately 100 to 392 ms after pronoun onset (schwa) and from approximately 198 to 392 ms (asterisk), across all scalp sites; whereas only *-\** differed from the incongruent condition -*a* showing a significant negative cluster (mass = −8886.692, *p* = .009) from approximately 342 to 666 ms across all sites. (ii) For stereotypically gender-congruent words (e.g., *chirurga*/*o*, “female/male surgeon”), both *-ə* and *-\** conditions did not differ significantly from the congruent condition -*o* (*p*s > .10), while they both differed from the incongruent condition yielding negative clusters (schwa: mass = −11542.475, *p* = 0; asterisk: mass = −12602.350, *p* = .002) spanning approximately 350 to 800 ms across all sites. (iii) For stereotypically gender-incongruent words (e.g., *maestra/o*, “female/male teacher”) both *-ə* and *-\** conditions differed significantly from the congruent condition *-o* yielding negative clusters (schwa: mass = −6812.389, *p* = .012; asterisk: mass = −6411.978, *p* = .011), with effects spanning approximately 100–456 ms for schwa (centro-parietal sites) and 100–372 ms for asterisk (all sites); and from the incongruent condition -*a*, yielding negative clusters (schwa: mass = −22847.816, *p* = 0; asterisk: mass = −18681.056, *p* = .0008), extending across the full time window at centro-parietal sites (schwa) and from approximately 316 to 800 ms across all sites (asterisk). Responses to *lui* did not differ between schwa and asterisk conditions (*p*s > .10). This pattern suggests that the masculine pronoun is ultimately integrated into gender-inclusive representations no matter the word stereotype content (as represented by the lack of late positivity), while initial integration difficulty (N100-N400 effects) is absent for stereotypically congruent words.

In response to the pronoun *lei*, (i) for stereotype-neutral words (e.g., *vicina/o*, “female/male neighbor”), neither *-ə* nor *-\** conditions differed significantly from the congruent *-a* nor the incongruent -*o* conditions (*p*s > .08). (ii) For stereotypically gender-congruent words (e.g., *maestra/o*, “female/male teacher”), neither *-ə* nor *-\** conditions differed significantly from the congruent condition -*a* (*p*s > .10), but *-\** differed significantly from the incongruent condition *-o*, yielding a negative cluster (mass = −4520.868, *p* = .031) extending from approximately 420 to 626 ms at all scalp sites. (ii) For stereotypically gender-incongruent words (e.g., *chirurga*/*o*, “female/male surgeon”), neither *-ə* nor *-\** conditions differed significantly from the congruent condition -*a* (*p*s > .06), while *-ə* differed significantly from the incongruent condition *-o*, yielded two negative clusters, the earliest from approximately 120 to 220 ms after pronoun onset (mass = −3430.227, *p* = .031), the latest from approximately 388 to 558 ms (mass = −3677.130, *p* = 0.029), both across all sites. Responses to *lei* did not differ between schwa and asterisk conditions (*p*s > .10). This pattern suggests that, as for the masculine pronoun, the feminine pronoun is ultimately integrated into gender-inclusive representations (as represented by the lack of late positivity) at least for schwa forms for stereotypically gender-congruent words, and asterisk for stereotypically gender-incongruent words.

### No Stereotypical Gender ERP Priming Effect for Any Ending Form

Last, we examined the EEG response to pronouns primed by stereotypically male and stereotypically female words, within each ending form (i.e., *-o, -a, -ə*, *-\**) (see Table 2). Our goal was to assess whether neutralization strategies effectively reduce also *stereotypically* triggered gender expectations.

We found no gender stereotype priming effects (i.e., differences between male-oriented vs female-oriented) for either masculine or feminine pronouns, across any ending category (all *p*s > .06). To better understand these null findings and directly assess the stereotypical gender expectations elicited by our prime stimuli irrespective of the endings, we conducted a follow-up behavioral study in which 31 participants (*M*_age_ = 22.48, *SD* = 2.67) gender-categorized the masculine and feminine Italian pronouns, primed by the word roots of our stimuli, as well as the common-gender nouns used in Siyanova-Chanturia et al. (2012). The details of the present follow-up are reported in the Supplementary Material.

We fitted a linear mixed-effect model with RTs to the target as dependent variable, Target Gender (masculine, feminine), Prime Stereotype (male-oriented, female-oriented, none) and Prime Type (root, common gender noun) as predictors along with their three-way interaction and participants and items as random intercepts.

The model showed a significant Target Gender × Prime Stereotype interaction, indicating that the feminine pronoun *lei* was responded faster when preceded by feminine than masculine primes, whereas the opposite pattern emerged for *lui*. Importantly, this effect did not vary as a function of Prime Type, as the three-way interaction was not significant, suggesting that the stereotype effect was consistent across both word roots and traditional common-gender nouns.

## Discussion

The present study aimed to provide neural evidence on whether GIL forms can neutralize grammatical gender cues and also attenuate gender stereotypes. This evidence directly informs the intense public, political, and academic debate on GIL efficacy and costs, by showing how these novel linguistic forms are processed in real time. We adopted a priming paradigm, well suited to compare the activation of implicit/automatic gender expectations elicited by gender inclusive and canonical forms. Italian speaking participants were presented with with nouns carrying either canonical (masculine *-o*, feminine *-a*) or neutralized (*-*, *-ə*) endings, varying in stereotypical association (none, e.g., *vicina/o/ǝ/\**, “neighbor”; male-oriented, e.g., *chirurgo/a/ǝ/\**, “surgeon”; female-oriented, e.g., *maestra/o/ǝ/\**, “teacher”), followed by masculine or feminine personal pronouns (*lui*, “he”; *lei*, “she”), which they categorized by gender. By recording RTs and ERPs to target pronouns, we asked: first, whether inclusive forms support gender-neutral representations (indexed by equal integration of masculine and feminine targets); and second, whether they also modulate stereotype-driven influences on pronoun processing.

### Neutralization Strategies Attenuate Morphology-Driven Gender Expectations

Canonical (masculine *-o*, feminine *-a*) word endings produced strong implicit and automatic gender expectations, so that, as predicted, participants were faster to gender-categorize the masculine pronoun *lui* (“he”) after masculine-marked words (*-o*), and *lei* (“she”) after feminine ones (*-a*). Accompanying ERP responses signaled lexical-semantic integration difficulties and morphosyntactic reanalysis for incongruent pronouns, expressed as an P300/LPP effect for *lui* after*-a,* and a biphasic N400-P300 effect for *lei* after *-o* (Casado et al., 2023; Chalyvidou & Weber, 2025; Pesciarelli et al., 2019; Serafini & Pesciarelli, 2025, 2026; Siyanova-Chanturia et al., 2012). A slightly different effect for the two pronouns is consistent with emerging evidence for a neural gender asymmetry, predicting that feminine forms elicit stronger gender expectancy constraints than the masculine one (Alemán Bañón & Rothman, 2016; Beatty-Martínez & Dussias, 2019; Beatty-Martínez et al., 2021).

Gender-inclusive endings (-*\**, -*ǝ*), by contrast, produced weaker implicit and automatic gender expectations. Participants’ speed to gender-categorize pronouns following these forms fell exactly between that for congruent and incongruent pronouns, with no difference between the two neutralized forms. In other words, neither pronoun was expected or unexpected after GIL, as predicted if it carried gender-neutral cues. Accompanying ERP responses showed that both pronouns following GIL forms elicited early differences from congruent and later differences from the incongruent pronouns, suggesting initial processing difficulties followed by successful integration of both genders.

The initial processing difficulties likely reflect increased lexical-semantic integration demands for pronouns following GIL forms relative to gender-matching canonical forms, as indexed by the larger N400. This may arise because GIL forms provide weaker retrieval cues from semantic memory (Kutas & Federmeir, 2000; Lau et al., 2008). Alternatively, although the pronoun itself is readily processed, integrating its meaning with the preceding gender-inclusive context may be more demanding (Friederici, 2002; Hagoort, 2008). Notably, the schwa-*lei* condition did not produce this effect, suggesting *-ǝ* may prime feminine targets similarly to *-a* endings, likely due to graphovisual similarity— though no overall difference between schwa and asterisk was found.

In addition to an expected N400 effect, an unpredicted N100-like effect emerged selectively for masculine pronouns following GIL relative to canonical endings. This effect may reflect increased attentional engagement with a less expected masculine continuation following GIL, potentially an early female bias, in line with the N100 association with perceptual and attentional processes (Herrmann & Knight, 2001), and its sensitivity to emotional relative to neutral words (Bernat et al., 2001; Díaz-Lago et al., 2015; Fraga et al., 2021; Hofmann et al., 2009; Kissler et al., 2006; Scott et al., 2009).

Crucially, neither pronoun following GIL forms triggered morphosyntactic reanalysis, repair processes, or context updating mechanisms indexed by P300/P600 effects (Donchin & Coles, 1988; Friederici, 2011). This is the strongest evidence that neutralized forms can elicit gender-balanced representations, since neither pronoun after GIL represents a grammatical gender violation.

This neural time course for GIL in Italian closely aligns with recent Spanish behavioral evidence: Román Irizarry et al. (2025) showed that innovative endings such as *-e* and *-x* (e.g., *todes*, *todxs*, “all”) generate processing costs relative to canonical binary forms, but without selectively privileging masculine or feminine interpretations. Similarly, psycholinguistic studies in Spanish reported that non-binary forms are processed more slowly than canonical morphology while remaining interpretable as genuinely inclusive forms (Stetie & Zunino, 2022; Zarwanitzer & Gelormini-Lezama, 2023).

This findings however contrasts with the only prior ERP study examining neo-morpheme processing, i.e., the German gender star (e.g., *Lehrer*innen*, “teachers”) (Glim et al., 2025), which reported a P600 for male continuations, interpreted as evidence of a female-biased representation. It similarly contrasts with prior behavioral literature identifying female biases, i.e., increased cognitive availability of women (Abbondanza et al., 2025; Glim et al., 2025; Körner et al., 2022; Pozniak et al., 2024; Tibblin et al., 2023). These cross-linguistic differences are informative: forms like the German gender star and French midpoint (e.g., *citoyen·ne·s*, “citizens”) explicitly combine masculine and feminine morphology, potentially increasing feminine salience, whereas Italian and Spanish inclusive forms attempt to neutralize binary morphology altogether. Regarding the findings of Abbondanza et al. (2025), it is important to note that their study relied on explicit gender-association ratings. As participants consciously evaluated the gender associated with neutralized forms, the observed female bias may reflect pragmatic interpretations of the communicative goals of inclusive language—namely, increasing the visibility of women—rather than automatic gender representations during online language processing.

Importantly, most prior studies embedded inclusive forms in sentences or discourse, where multiple syntactic, semantic, and pragmatic cues contribute to interpretation. Here, using a priming paradigm with isolated word, targeting rapid and automatic gender expectations, the neutralized endings were the primary source of gender information available. The fact that both *-ǝ* and *-\** yielded intermediate RTs for both pronouns and no P300 response therefore provides particularly strong evidence that these forms attenuate automatic gender predictions rather than biasing interpretation toward either gender. Figure 5 illustrates the proposed time course of gender processing following canonical and gender-inclusive word endings.

**Figure 5.**
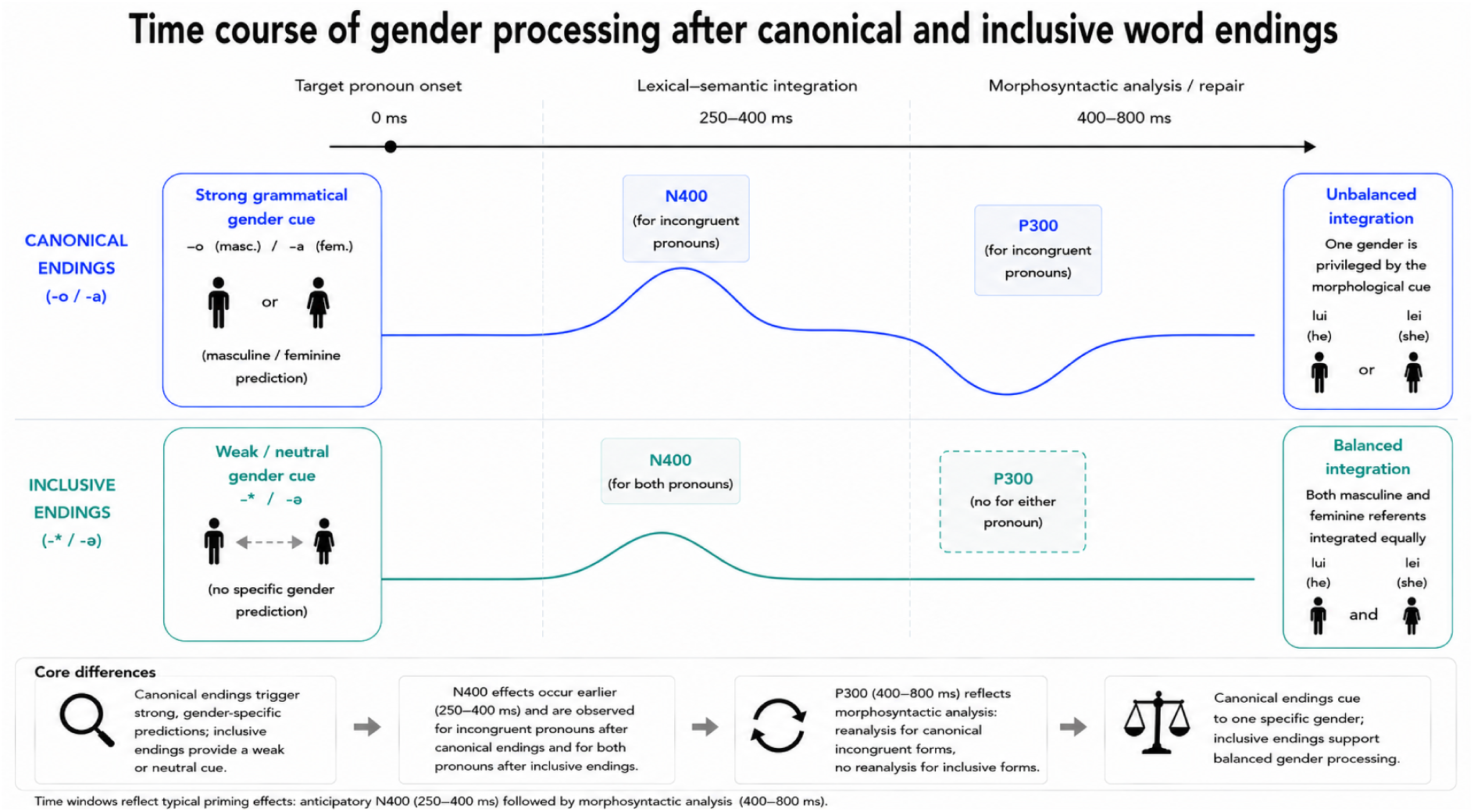
Proposed time course of gender processing following canonical and gender-inclusive word endings. Canonical word endings provide a strong grammatical gender cue, eliciting an N400 effect for gender-incongruent pronouns followed by a late P300 response, consistent with morphosyntactic reanalysis or repair. By contrast, gender-inclusive endings elicit an N400 effect for both pronouns, with no subsequent P300, suggesting successful lexical-semantic integration without reanalysis. Thus, canonical endings promote unbalanced integration by favoring one gender representation over the other, whereas gender-inclusive endings support balanced integration of masculine and feminine referents. The schematic ERP waveforms summarize the qualitative pattern of effects observed in the present study and are intended for illustrative purposes only.

### Stereotype-Driven Expectations Appear to Influence Initial Processing Costs but Less so Later Integration Mostly for Masculine Pronouns

An important question is whether gender-inclusive forms merely reduce morphologically driven biases or also attenuate stereotype-based gender representations. We first asked whether the effects so far reported varied as a function of the stereotypical association of words. Second, we asked whether GIL forms, which act on word endings but leave the word stem unchanged, left any residual stereotype effects unresolved.

Answering the first question, contrary to predictions, behaviorally stereotype category did not modulate grammatical or GIL effects, with no interactions between stereotype category, prime ending, and target gender in RTs. At the neural level, also unpredicted, processing difficulties were not selectively stronger when morphology and stereotypes jointly mismatched the pronoun (e.g., avvocato “male lawyer” – *lei* “she”), but they were equally present even when only morphology mismatched (e.g., maestro “male teacher” – *lei* “she”); and, second, they were absent for stereotype-neutral words (e.g., vicino “male neighbor” – *lei* “she”). This contrasts with previous studies (Pesciarelli et al., 2019; Siyanova-Chanturia et al., 2012), in which gender-marked words were specifically selected to be stereotype-neutral, and stereotypical words to be grammatically gender neutral, i.e., common gender (e.g., *conducent**e*** “driver”). Here, the combination of overt gender-marking and stereotypical association may have increased lexical salience, thereby amplifying grammatical gender effects.

For GIL forms, early processing costs for the masculine pronoun emerged only after stereotypically neutral or stereotypically incongruent primes (e.g., *vicin\** “neighbor” – *lui* “he”; *maestr\** “teacher” – *lui* “he”), but not after stereotypically congruent ones (e.g., *avvocat\** “lawyer” – *lui* “he”), suggesting that stereotypical congruency facilitated lexical-semantic integration of masculine continuations, potentially biasing GIL representations toward male referents. The additional N100-like effect emerged only for masculine pronouns after stereotypically incongruent GIL forms (e.g., *maestr*/ǝ* “teacher”– *lui* “he”) relative to canonical forms (e.g., *maestro/a* “male/female teacher”– *lui* “he”), suggesting that the increased attentional allocation to unexpected continuations is mostly driven by stereotypical violations. No corresponding processing costs were observed for feminine pronouns, suggesting that these are more evident for masculine pronouns overall, in line with evidence for a female bias in GIL processing, and that stereotypical information exerts stronger early influence on masculine continuations (Serafini & Pesciarelli, 2025, 2026)—either facilitating integration when congruent or imposing additional costs when incongruent.

Despite these early modulations, masculine continuations were ultimately integrated into GIL representations regardless of stereotype category, thus even when violating stereotypical expectations (e.g., *maestr\** – *lui* “teacher-he”). The exception was the schwa condition, which did not show later integration for stereotype-neutral primes (e.g., *vicinǝ – lui* “neighbor-he”, not different from *vicina – lui* “female neighbor-he”). For feminine pronouns, later integration was more selective, emerging only for stereotypically congruent asterisk forms (e.g., *maestr\** – *lei* “teacher-she”) and stereotypically incongruent schwa forms (e.g., *chirurgǝ – lei* “surgeon-she”), with schwa possibly more effective at integrating feminine pronouns when the stereotype mismatches, given its morphological similarity to *-a*. The relative absence of stereotype modulation at the later stage is compatible with evidence that gender stereotypes primarily impact processing at the N400 stage (Pesciarelli et al., 2019; Siyanova-Chanturia et al., 2012), mostly in case of masculine violations (Serafini & Pesciarelli, 2025, 2026). Importantly, they do not prevent later integration.

Answering the second question, RTs and ERPs to both pronouns did not differ between stereotypically congruent and incongruent condition for either canonical (e.g., *chirurgo* – *lui* “male surgeon-he” vs. *maestro – lui* “male teacher-he”) or GIL endings (e.g., *chirurg\** – *lui* “surgeon-he” vs. *maestr* – lui* “teacher-he”). This absence of stereotype-congruency effects contrasts with our follow-up study, in which the same stems, presented without endings, elicited stereotypical congruency effects comparable to those for common-gender nouns used in previous studies (Pesciarelli et al., 2019; Siyanova-Chanturia et al., 2012), suggesting the null effects here cannot be attributed to the stimuli themselves.

Grammatical gender may have overridden stereotypical gender during online processing: stereotypically incongruent pairs such as *chirurga – lei* “female surgeon- she” or *maestro – lui* “male teacher-he” may have been processed as fully congruent due to morphosyntactic match, as previously reported (e.g., *la presidente*, “the_[G,F]_ president” - female face; Navarrete et al., 2026; e.g., *esthéticiens*_[G,M]_ “beauticians”- some of the men; Gygax et al., 2008; but see: Carreiras et al., 1996; Molinaro et al., 2016). Notably, GIL forms should minimize grammatical gender marking, so the absence of stereotype effects here may indicate that these forms reduce not only grammatical gender biases but also stereotypical gender expectations.

An alternative interpretation is that participants may have strategically allocated attention to word endings, the critical manipulation, at the expense of the lexical stem, where stereotypical meaning is encoded, thereby attenuating activation of stereotypical gender knowledge. Further work should disentangle these accounts.

Fewer studies on GIL have examined stereotype-associated words. Our findings show that gender stereotypical expectations mostly influence attention to and lexical-semantic integration of (masculine) pronouns, allowing the successful integration of both pronouns into GIL-elicited representations. This findings align with Stetie and Zunino’s (2024) evidence of no effect of stereotypical association strength, high (e.g., *lxs plomerxs* “the plumbers”) compared to low (e.g., *lxs maestrxs* “the teachers”), for Spanish *-x/-e* forms, which overall elicited more mixed-gender representations without additional processing costs relative to the generic masculine. Stronger stereotypical effects were instead obtained by Xiao et al. (2023), who found that the French mid-dot form produced an overestimation of the proportion of women for male-stereotyped professions (e.g., *électricien·ne* “electrician”), an overestimation of men for female-stereotyped professions (e.g., *caissier·ère* “cashier”), while consistent estimates for non-stereotyped ones.

Together, these findings suggest that gender-inclusive forms could interact with pre-existing social knowledge tied to specific roles. Crucially, the absence of differential facilitation for masculine versus feminine pronouns and the integration at the P300 stage in the present study indicates that, although stereotypical knowledge remained available during processing, neutralized forms did not bias interpretation toward either gender. These findings suggest that such forms may serve as viable candidates for gender-neutral reference, with broader implications for inclusivity in language use.

## Conclusions

Our findings provide the first ERP evidence that gender-inclusive endings in Italian attenuate automatic grammatical gender predictions without systematically favoring either masculine or feminine representations. While inclusive forms incurred an early processing cost relative to canonical congruent forms, they did not elicit the later morphosyntactic reanalysis typically associated with gender violations, indicating that both masculine and feminine continuations can ultimately be integrated within a neutral representation. These findings provide a neurocognitive account of how emerging inclusive forms are processed and offer a mechanistic framework for understanding how linguistic innovations may reshape automatic gender representations during language comprehension. More broadly, they establish an experimentally controlled and adaptable paradigm for investigating the online processing of current and future gender-inclusive innovations across languages.

## Resource Availability

### Lead Contact

Requests for further information and resources should be directed to and will be fulfilled by the lead contact, Luana Serafini.

### Materials Availability

This study’s material has been deposited to OSF at: https://doi.org/10.17605/OSF.IO/EPGUH

### Data and Code Availability

Behavioral and ERP data have been deposited at OSF and are publicly available as of the date of publication at: https://doi.org/10.17605/OSF.IO/EPGUH

All original code has been deposited at OSF and is publicly available at https://doi.org/10.17605/OSF.IO/EPGUH as of the date of publication.

Any additional information required to reanalyze the data reported in this paper is available from the lead contact upon request.

## Acknowledgments

We are grateful to Riccardo Zavalloni, Chiara Maria Salerno and Alessandra Cristiano for assistance with participant’s recruitment and data collection. The study was funded by the PNRR - Missione 4 ‘Istruzione e Ricerca’ - Componente C2 Investimento 1.1 ‘Fondo per il Programma Nazionale di Ricerca e Progetti di Rilevante Interesse Nazionale (PRIN)’, ‘Gender bias in language: Testing INClusive ITalian language feasibility and impact (INCITƏ)’, codice progetto 202272BY39 (CUP E53D23011740006). Funding sources had no involvement in the study design, collection, analysis and interpretation of data, writing of the report and decision to submit the article for publication.

## Author Contribution

L.S., M.A., F.P.: Conceptualization, Methodology, Validation, Project administration. L.S., M.A.: Software, Formal analysis, Investigation, Data curation, Visualization, Writing- Original draft preparation. F.P.: Resources, Supervision, Funding acquisition, Writing- Reviewing and Editing.

## Declaration of Interests

The authors declare no competing interests.

## Supplemental Information

**Document S1.** Supplemental methods, Tables S1 and S2, supplemental references.

## Notes

https://doi.org/10.17605/OSF.IO/EPGUH

### Competing Interest Statement

The authors have declared no competing interest.

