## Supplementary Material for "Neural Evidence That Gender-Inclusive Language Attenuates Automatic Gender Predictions"

This manuscript is a preprint and has not been peer-reviewed.

This PDF file includes:
Supplementary Methods
Table S1 and S2
Supplemental references

**Norming study for prime words**

A set of 305 gender-marked words (including nouns, past participles, and adjectives referring to occupations, roles, or personal characteristics) ending with the canonical Italian nominal endings –*a*[F]/-*o*[M] were included in an online questionnaire administered to 41 participants (5 men, $M_{age}$ = 22,53 yrs; age range 20-34 yrs) not further involved in the experiment.

Participants rated on a 9-point Likert scale the extent to which each word was associated with a masculine or feminine stereotype. The instructions were as follows: "*You will find below a list of words accompanied by a 9-point rating scale. Please indicate whether the word is generally associated with a male or a female stereotype. The scale ranges from 1 to 9, where 1 means exclusively associated with a male stereotype and 9 exclusively associated with a female stereotype. A score of 5 indicates that the concept refers equally to both a male and a female stereotype.*" Scale labels (1 – only men, 5 – both, 9 – only women) were reversed for half of the participants. The final rating assigned to each word was calculated by combining the ratings obtained with both directions of the scale. In addition, half of the participants evaluated the words presented in the masculine/feminine form (e.g., *maestro/a*, "male/female teacher"), while the other half evaluated the words presented in the feminine/masculine form (e.g., *maestra/o*, "female/male teacher"), in order to avoid biases due to the presentation order of the ending.

From all rated words, we selected the stimuli as belonging to three categories: (i) 30 stereotype-neutral words (ranging from 4.7 to 5.3 points), (ii) 30 stereotypically male words (scoring < 3.5 points), (iii) 30 stereotypically female words (scoring > 6.5 points).

Twenty filler words were also included. These common gender words (i.e., ending in -*e* or consonant, as for loans) were selected from a different questionnaire. A set of 641 words (including nouns, past participles, and adjectives referring to occupations, roles, or personal characteristics) was presented in three online questionnaires, containing 214, 214 and 213 items, respectively, and administered to a total of 123 university students and researchers' acquaintances (80 women, 1 non-binary, $M_{age}$ = 23.36 yrs, $SD_{age}$ = 6.33) who did not take part in the main experiment. All words were in common gender form, meaning they could refer to both male and female individuals. Participants were asked to rate the extent to which each word was associated with men, women, or both, using a seven-point Likert scale (1 = only men, 4 = both, 7 = only women). Whenever participants did not know the meaning of the word, they could opt for the "I don't know" choice. To control for potential response biases, the direction of the scale was reversed for half of the participants. For each word, a final stereotypicality score was calculated by averaging ratings across both scale orientations.

The 20 words selected as filler primes received ratings between 3.8 and 4.2, indicating no stereotypical association.

**Table S1**. *Properties and balancing of prime stimuli across stereotype categories*

| | Stereotypically neutral | Stereotypically masculine | Stereotypically feminine | |
|---|---|---|---|---|
| Frequency (Zipf) | 3.39 | 2.92 | 3.06 | .091 |
| Length (characters) | 7.96 | 8.36 | 8.20 | .654 |
| Valence | 4.29 | 4.04 | 4.01 | .067 |
| Stereotypicality (absolute value) | / | 1.65 | 1.57 | .431 |

**Note.** P-values refer to one-way ANOVAs testing differences across stereotype categories. Stereotypicality was not applicable (/) for the neutral category.

**Individual measures' impact on response times**

We tested whether the variables collected in the post-experimental questionnaires (cfr. Section "Design and Procedure") modulated the Prime Ending × Target Gender interaction running a full model containing three three-way interactions (e.g. Prime Ending × Target Gender × Measure). The model included fixed effects of Prime ending, Target, participant-level covariates (Feeling LGBT, Ambivalent Sexism Inventory [ASI], Bem femininity, Bem masculinity, Knowledge, and Attitude), their interactions with Prime ending and Target, and Prime gender stereotype. Random intercepts were included for participants (n = 36) and items (n = 90). Continuous predictors were mean-centered. Estimates (β), standard errors (SE), degrees of freedom (df), *t*-values, and *p*-values are reported.

The questionnaire measures included ASI, BSRI, and Feeling Thermometer, familiarity with gender-inclusive language, and perceived usefulness and applicability of gender-inclusive language. Below, we report the significant results that emerged from the model, see the full output in Table S2.

The Prime Ending × Target Gender × Feeling Thermometer interaction was significant for the *-o* and *asterisk* conditions when TARGET was *lei* (Est. = .0007, $t$ = 2.23, $p$ = .025; Est. = .0008, $t$ = 2.46, $p$ = .013), indicating that higher LGBT feeling scores were associated with shorter response times for *lei* in the *-a* and schwa conditions. For *lei*, the Prime Ending × Target Gender × ASI interaction was significant for the *-o*, asterisk, and schwa conditions (Est. = -0.0614, $t$ = -4.40, $p$ < .001; Est. = -0.0557, $t$ = -3.99, $p$ < .001; Est. = -0.0446, $t$ = -3.19, $p$ = .001), indicating that higher ASI scores were associated with longer response times, but the pattern was substantially reduced for *lui*. The Prime Ending × Target

Gender × BEM interaction was significant for *lei* in the -o and asterisk conditions (Est. = -0.0893, $t$ = -4.91, $p$ < .001; Est. = -0.0400, $t$ = -2.20, $p$ = .028), whereas the effect for schwa was not significant (Est. = -0.0204, $t$ = -1.12, $p$ = .263), indicating that higher femininity scores were associated with shorter response times for *lei* mainly in the o and asterisk conditions.

**Table S2.** *Fixed effects from the linear mixed-effects model predicting log-transformed response times.*

| Predictor | Estimate (β) | SE | df | *t* | *p* |
|---|---|---|---|---|---|
| (Intercept) | 6.033 | 0.153 | 30.33 | 39.357 | < .001 |
| prime o × TARGET × Feeling LGBT | 0.00074 | 0.00033 | 24470 | 2.230 | .026 |
| prime -ast × TARGET × Feeling LGBT | 0.00081 | 0.00033 | 24470 | 2.466 | .014 |
| prime schwa × TARGET × Feeling LGBT | 0.00059 | 0.00033 | 24470 | 1.786 | .074 |
| prime o × TARGET × ASI | -0.061 | 0.014 | 24470 | -4.404 | < .001 |
| prime -ast × TARGET × ASI | -0.056 | 0.014 | 24470 | -3.994 | < .001 |
| prime schwa × TARGET × ASI | -0.045 | 0.014 | 24470 | -3.187 | .001 |
| prime o × TARGET × Bem femininity | -0.089 | 0.018 | 24470 | -4.906 | < .001 |
| prime -ast × TARGET × Bem femininity | -0.040 | 0.018 | 24470 | -2.198 | .028 |
| prime schwa × TARGET × Bem femininity | -0.020 | 0.018 | 24470 | -1.120 | .263 |
| prime o × TARGET × Bem masculinity | 0.009 | 0.011 | 24470 | 0.796 | .426 |
| prime -ast × TARGET × Bem masculinity | 0.014 | 0.011 | 24470 | 1.287 | .198 |
| prime schwa × TARGET × Bem masculinity | 0.009 | 0.011 | 24470 | 0.792 | .428 |
| prime o × TARGET × Knowledge | -0.001 | 0.006 | 24470 | -0.225 | .822 |
| prime -ast × TARGET × Knowledge | -0.004 | 0.006 | 24470 | -0.669 | .504 |
| prime schwa × TARGET × Knowledge | -0.002 | 0.006 | 24470 | -0.410 | .682 |
| prime o × TARGET × Attitude | -0.012 | 0.007 | 24470 | -1.856 | .063 |
| prime -ast × TARGET × Attitude | -0.011 | 0.007 | 24470 | -1.600 | .110 |
| prime schwa × TARGET × Attitude | -0.005 | 0.007 | 24470 | -0.744 | .457 |

**Note.** Linear mixed-effects model predicting log-transformed response times. Random intercepts were included for participants ($n$ = 36) and items. Continuous predictors were mean-centered. Estimates (β), standard errors (SE), degrees of freedom (df), $t$-values, and $p$-values are reported.

**Follow-up study**

**Participants**

Thirty-two students (27 female; $M_{age}$ = 20.37, $SD_{age}$ = 1.11) from the University of Modena and Reggio Emilia took part in the study for course credit. None had participated in the main experiment. All were Italian monolingual speakers with no reported history of language or neurological disorders and normal or corrected-to-normal vision. Written informed consent was obtained from all participants.

**Materials**

A total of 180 prime stimuli were created for the follow-up study. Primes varied along two dimensions: word ending (common-gender vs. stem-only) and gender stereotype (masculine, feminine, neutral), yielding a balanced design with 30 items per category.

The stem-only primes were derived from the same prime words used in the main experiment by removing the final vowel encoding grammatical gender (*-a*/*-o*), leaving the bare lexical stem (e.g., *chirurg-*, "surgeon"). Because these stems carry no morphological gender marking, they allowed us to test whether our stimuli were effective in automatically activating gender stereotypes in the absence of any grammatical gender cue. The common-gender primes, by contrast, consisted of word forms that are inherently unmarked for grammatical gender (e.g., *conducente*, "driver"). These same stimuli have been used in prior priming studies that reliably reported stereotypical priming effects (Siyanova-Chanturia et al., 2012; Pesciarelli et al., 2019; Serafini & Pesciarelli, 2025, 2026), and served here as a benchmark condition against which the stem-only primes could be compared.

**Procedure**

Participants first completed 12 practice trials, followed by 360 experimental trials divided into three blocks. On each trial, a prime word was presented before a target pronoun (*lui* “he” or *lei* “she”) and participants had to indicate the gender of the target pronoun by pressing a key on the keyboard. Key assignment was counterbalanced across participants. Response times and response accuracy were recorded for each trial.

**Statistical analysis & results**

Response times (RTs) were log-transformed prior to analysis. We fitted a linear mixed-effects model using the lme4 package in R, with Target Gender (*lui* “he” vs. *lei* “she”), Prime Gender Stereotype (female vs. male), Prime Type (common-gender vs. stem only), and all interactions as fixed effects. Random intercepts were included for participants and items.

The analysis revealed that the three-way interaction among Target Gender, Prime Gender Stereotype, and Prime Type was not significant ($t$ = 1.49, $p$ = .136); while the interaction between Target and Prime Gender Stereotype was ($t$ = −4.53, $p$ < .001).

Follow-up pairwise comparisons showed that for the feminine target *lei*, responses were significantly faster following feminine than masculine primes ($z$ = −4.08, $p$ < .001). Conversely, for the masculine target *lui*, responses were significantly faster following masculine than feminine primes ($z$ = 2.61, $p$ = .009). These findings indicate a congruency effect between the gender stereotype conveyed by the prime and the target pronoun, irrespective of prime type.
